# Automated Navigation of the lncRNA Transcriptome: A comprehensive SnakeMake based computational Pipeline for robust Identification of lncRNAs and their putative targets

**DOI:** 10.1101/2024.08.18.608522

**Authors:** Manu Kandpal, Chitranjan Mukherjee, Bhadresh Rami

## Abstract

**Background:** Long non-coding RNAs (lncRNAs) have emerged as potent regulatory elements in cellular processes. The substantial increase in transcriptomic data resulting from high-throughput RNA sequencing necessitates effective approaches for the identification and functional annotation of lncRNAs.

**Method:** To address this need, we have developed a SnakeMake-based pipeline. Our pipeline automates and integrates several key steps: 1) RNA-seq analysis using Hisat2 and stringTie, (2) lncRNA identification using inhouse python scripts and tools CPC2 and BLASTX, (3) prediction of *cis-* and *trans-* gene targets of lncRNAs, and (4) KEGG pathway enrichment to obtain biological insights. Importantly, the pipeline allows users to customize parameters for each step through a user-friendly configuration file (config.yaml), enhancing flexibility and ease of use. One of the distinctive features of our approach is its single command execution, facilitating multiple runs without the need for extensive user intervention. This not only enhances user convenience but also ensures reproducibility of analyses across different studies.

**Result:** We applied our pipeline on rice, sorghum, and human RNA-seq data, to identify (1) List of all differentially expressed transcripts., (2) List of differentially expressed lncRNAs, (3) lncRNA target genes, (4) Enriched pathways to which target genes belong and (5) Obtain a visualization output in the form of a bubble plot that depicts the enriched pathways. Our approach can help researchers obtain valuable biological insights into how lncRNAs contribute to various biological functions.

**Conclusion:** The distinctive features of our SnakeMake-based automation pipeline position it as a versatile asset for researchers seeking a user-friendly, adaptable, robust, and reproducible solution for pan species lncRNA analysis. By efficiently uncovering the regulatory roles of lncRNAs in cellular processes, this pipeline has the potential to shed light on various biological phenomena, such as developmental biology, disease progression, and cellular response to external stimuli.

**GRAPHICAL ABSTRACT:** This study presents a SnakeMake-based pipeline for identifying and annotating long non-coding RNAs (IncRNAs) from RNA sequencing data. It integrates RNA-seq analysis, IncRNA identification, gene target prediction, and pathway enrichment, with customizable parameters through a user-friendly configuration file. The pipeline’s single command execution enhances convenience and reproducibility. (The bubble chart in the figure is a representative chart and provided as an example.)

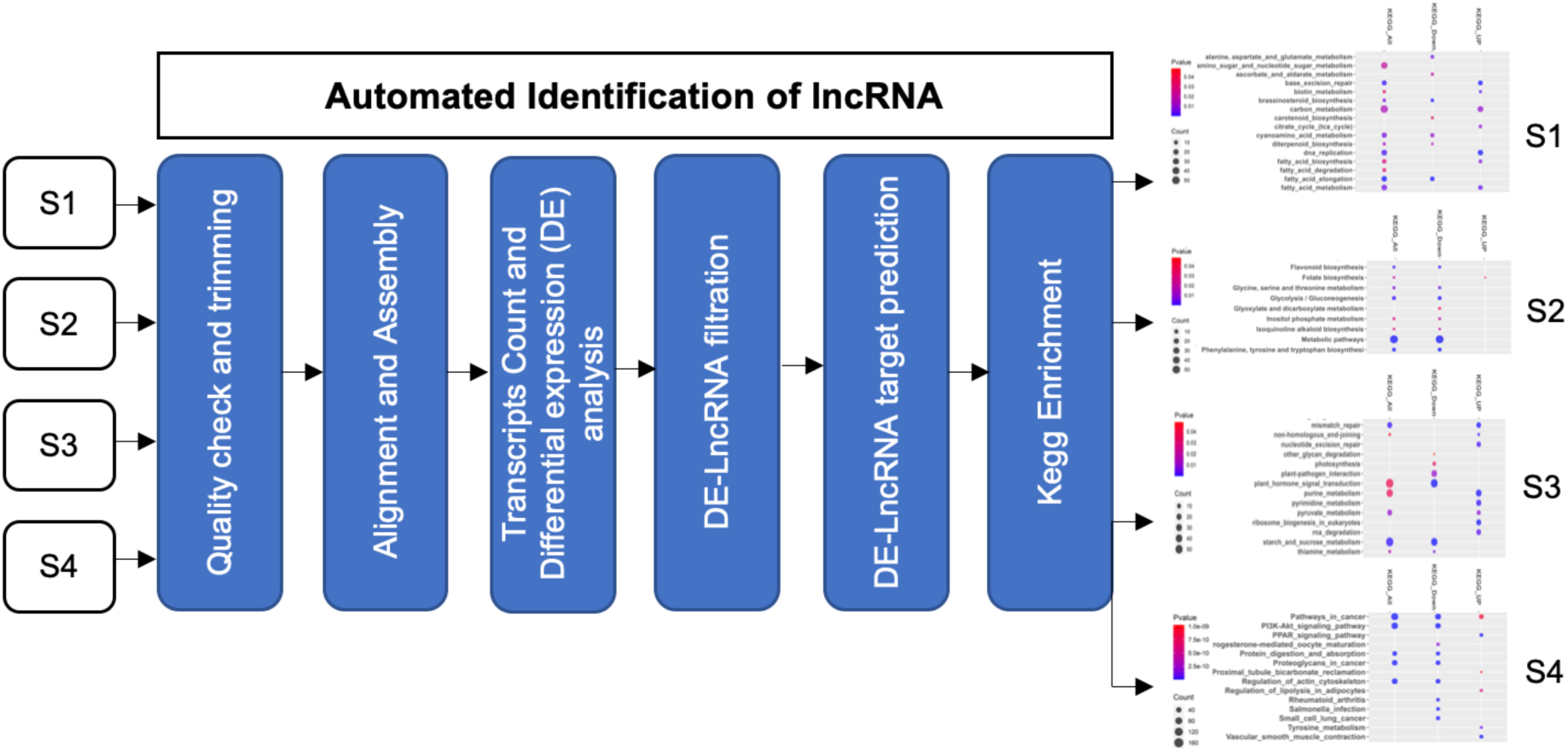

## 1. INTRODUCTION

Long noncoding RNAs (lncRNAs) constitute a diverse category of endogenous transcribed RNA that are characterized by transcript length of ≥ 200 nucleotides (Lan et al. 2021; Zhou et al. 2022). In contrast to protein-coding genes, lncRNAs do not code for proteins. However, their significance in the regulatory landscape of cellular processes has been increasingly recognized (Oo et al. 2022). lncRNAs can sequester miRNAs thus disrupting the binding on the target coding genes which ultimately results in indirect deregulation of the target genes of those miRNAs. lncRNAs can interact with transcription factors (Liu et al. 2018; Ai et al. 2019; Zhang et al. 2019), and chromatin modifier proteins (Chu et al. 2011; Li et al. 2017; Bell et al. 2018; Bonetti et al. 2020), thus regulating gene expression (Kitagawa et al. 2013; Xiang et al. 2014; Yang et al. 2015; Isoda et al. 2017; Saldaña-Meyer et al. 2019; Statello et al. 2021), apoptosis (Ma et al. 2014; Li et al. 2018; Tao et al. 2019), survival, cancer migration (Jiang et al. 2019; Wang et al. 2020; Ahmad et al. 2023), and metabolism (Lin et al. 2020; Liu et al. 2021; Agostini et al. 2022; Duan et al. 2023). Their involvement in these crucial pathways emphasizes the significance of lncRNAs in the overall framework of cellular regulation and disease (Schmitt and Chang 2016; Bartonicek et al. 2016; Bhan et al. 2017; Wang et al. 2023; Kohlmaier et al. 2023; Ahmad et al. 2023).

In plants, lncRNAs have been recognized for their roles in regulating development, stress responses, and other critical processes, and similarly, in humans they are involved in crucial biological processes and disease mechanisms. In plants, lncRNAs such as COOLAIR (Hawkes et al. 2016; Nguyen and Searle 2023) and COLDAIR (Kim et al. 2017) regulate flowering time through vernalization in Arabidopsis thaliana, while LDMAR influences male sterility in rice under long-day conditions (Ding et al. 2012a; Zhou et al. 2012; Ding et al. 2012b), and enod40 is essential for nodule formation during symbiotic nitrogen fixation in legumes (Gultyaev et al. 2023). MIEL1 and lncRNA39026 modulate plant immunity (Marino et al. 2013; Aznaourova et al. 2020) and stress responses (Hou et al. 2020; Zhang et al. 2023), respectively.

In humans, well-characterized lncRNAs include HOTAIR (Hajjari and Salavaty 2015), which is involved in gene silencing and cancer metastasis; MALAT1 (Arun et al. 2020), which regulates gene expression and alternative splicing; XIST (Li et al. 2022a), crucial for X-chromosome inactivation; and H19 (Liao et al. 2023), which plays roles in development and cancer. Other significant lncRNAs such as ANRIL (Aguilo et al. 2016), TERRA (Chebly et al. 2022), PVT1 (Li et al. 2022b), and NEAT1 (Dong et al. 2018) are associated with cancer, telomere maintenance, MYC (Shtivelman and Bishop 1989; Wu et al. 2023) regulation, and paraspeckle formation (Hirose et al. 2014), respectively. These examples illustrate the diverse and critical roles lncRNAs play in both plant and human biology, highlighting their importance in development, stress responses, defense mechanisms, and disease pathogenesis.

Despite their prevalence across all eukaryotes and their key roles in regulating cellular roles, precious little is known about lncRNA’s and many unanswered questions remain regarding their biology, including their origin, genomic organization, evolution, and specific roles. Although there have been attempts to unravel the complex functions of lncRNAs, a large fraction of these molecules remains unidentified, unclassifiable and poorly characterized. Thus, there is a vital need in the field to develop robust analysis pipelines to address this key unmet need.

Researchers have proposed multi-step procedures for identifying lncRNA that necessitate a comprehensive understanding of each stage. However, this leads to a decrease in the reproducibility of the process as different users may implement each stage differently. Therefore, it is critical to maximize the consistency and efficiency of these analysis methods, particularly when dealing with large datasets, and to make sure that all necessary files are in place to support reproducibility and transparency. Few tools have been developed to speed up the multi-step process, such as Linc2function (Ramakrishnaiah et al. 2023), ICAnnoLncRNA (Pronozin and Afonnikov 2023), and lncRNADetector (Shukla et al. 2021), to identify lncRNAs from the fasta sequences. However, there aren’t many pipelines that can identify lncRNAs from raw fastq files, which contain unprocessed raw data. It is important and preferable to begin data analysis with unprocessed raw files to minimize data loss. A few pipelines, such as CALINCA (Talyan et al. 2021) and UClncR (Sun et al. 2017), are available but they only analyze human data and cannot be used for any other species.

Our pipeline, built using the Snakemake workflow management system (Köster and Rahmann 2012), explicitly defines the individual steps of the analysis workflow by specifying software environments and their corresponding dependencies in terms of libraries required for the installation. This ensures that every time data is analyzed, it is done using the identical software versions and dependencies, thereby ensuring reproducibility and reducing human intervention.

Our pipeline encompasses multiple stages of analysis, starting from the handling of raw fastq data to analyzing transcriptomics data, identifying lncRNAs, classifying them, performing differential expression (DE) analysis between test vs control samples, and pinpointing the trans and cis target genes regulated by the DE-lncRNAs. Furthermore, our pipeline conducts KEGG enrichment analysis to uncover the enriched pathways governed by the genes targeted by the DE-lncRNAs.

An important feature of our pipeline is its versatility, as it can simultaneously process multiple samples across different species. This capability is achieved by providing the reference genome fasta and gtf/gff files specific to each species, enabling comprehensive analysis with just one command.

## 2. MATERIAL AND METHODS

Our pipeline takes raw, unprocessed, paired-ended sequence files (fastq.gz/.fq.gz) as input. Fig. 1 shows the files and folder structure in a manner in which the raw files (from either in-house experiments or databases like GEO or SRA), genome file and the annotation files need to be stored in the folder. The run folder structure is crucial, emphasizing a main folder containing Snakemake scripts and config files. Within this run folder, individual dataset folders containing raw paired-end sequence files are organized, along with the index files and gtf of the genome. There is no limit on the number of datasets that can be analysed simultaneously using our pipeline, provided suitable computational resources are available. Prerequisites include pre-run configuration of the user-edited config file, indexed reference genome (by hisat2-build) for each dataset, installation of Linux-based tools (fastQC, multiQC, fastp, bbmap, hisat2, stringtie2, gffread, gffcompare, CPC2, BLASTX, subread, and bedtools), and R packages (stringR, tidyverse, dplyr, tidyr, DESeq2, data.table, optparse, gtools, and ashr).

**Fig 1.**
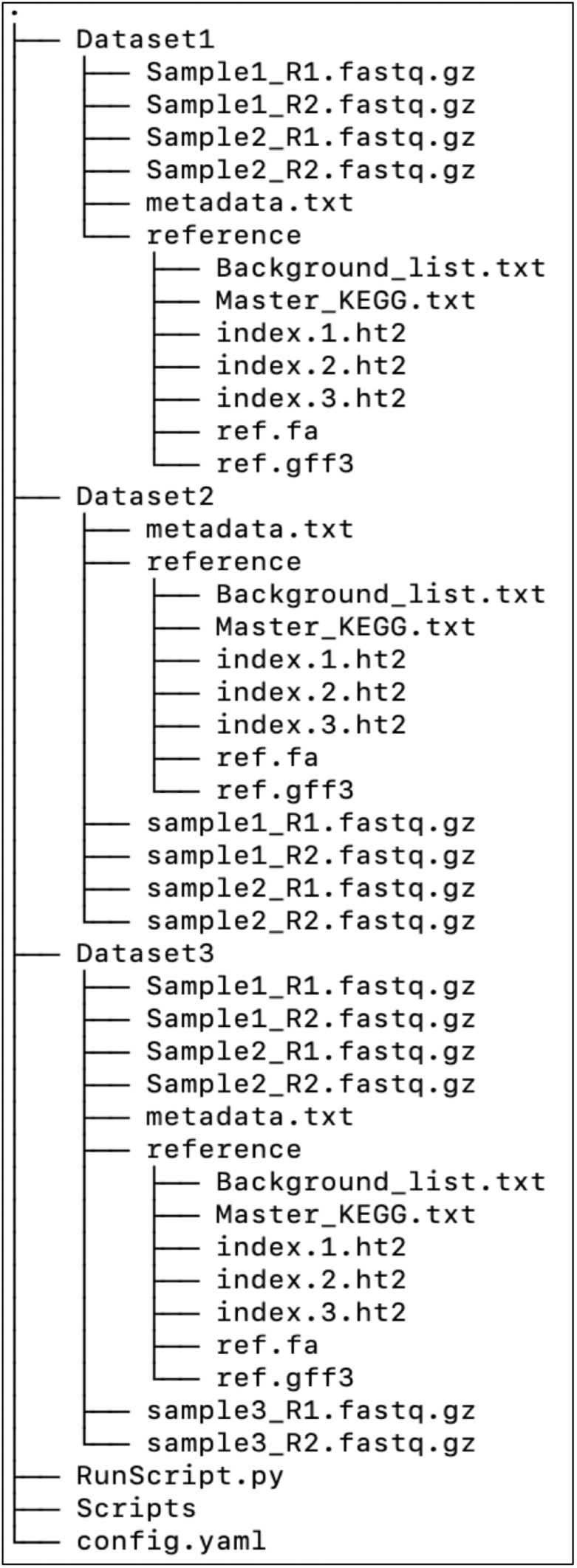
Structure of the working directory. Use this directory structure as a template to organize raw, genome, and metadata files in their respective folders. All result files generated during script execution will be saved in the dataset folder.

We have also provided a config.yaml file that can be edited by the users according to their requirements. For example, the following parameters can be modified by the user by easily editing the “config.yaml” file – (1) minimum length cut-off for trimming, (2) minimum phred score cut-off for trimming, (3) mapping percentage cut-off, (4) p-value and absolute log2FoldChange cut-off for differential expression analysis, (5) BLASTX E-value cut-off, (6) flanking length in BP upstream and downstream to the DE-lncRNA’s location, (7) p-value cut- off and Pearson correlation cut-off for identifying trans targets. The ability to modify the parameters by editing the config.yaml files provides significant flexibility to the individual user to finetune the pipeline to meet their study requirements as per experiment design. The details about each parameter is explained in the following sections.

Once the files are arranged in the proper folders, user is ready to run the pipeline for lncRNA identification developed in our study as shown in Fig 2. The pipeline is divided into two scripts. The first script performs quality filtering, transcript mapping to genome and transcriptome assembly, while the second script performs lncRNA identification, origin-based classification, DE analysis, identification of *cis-* & *trans-* targets genes of the DE- lncRNA’s, and KEGG pathway enrichment of these target genes. The stepwise details of the pipeline, as outlined below, provide user-friendly guidelines for execution.

**Fig 2.**
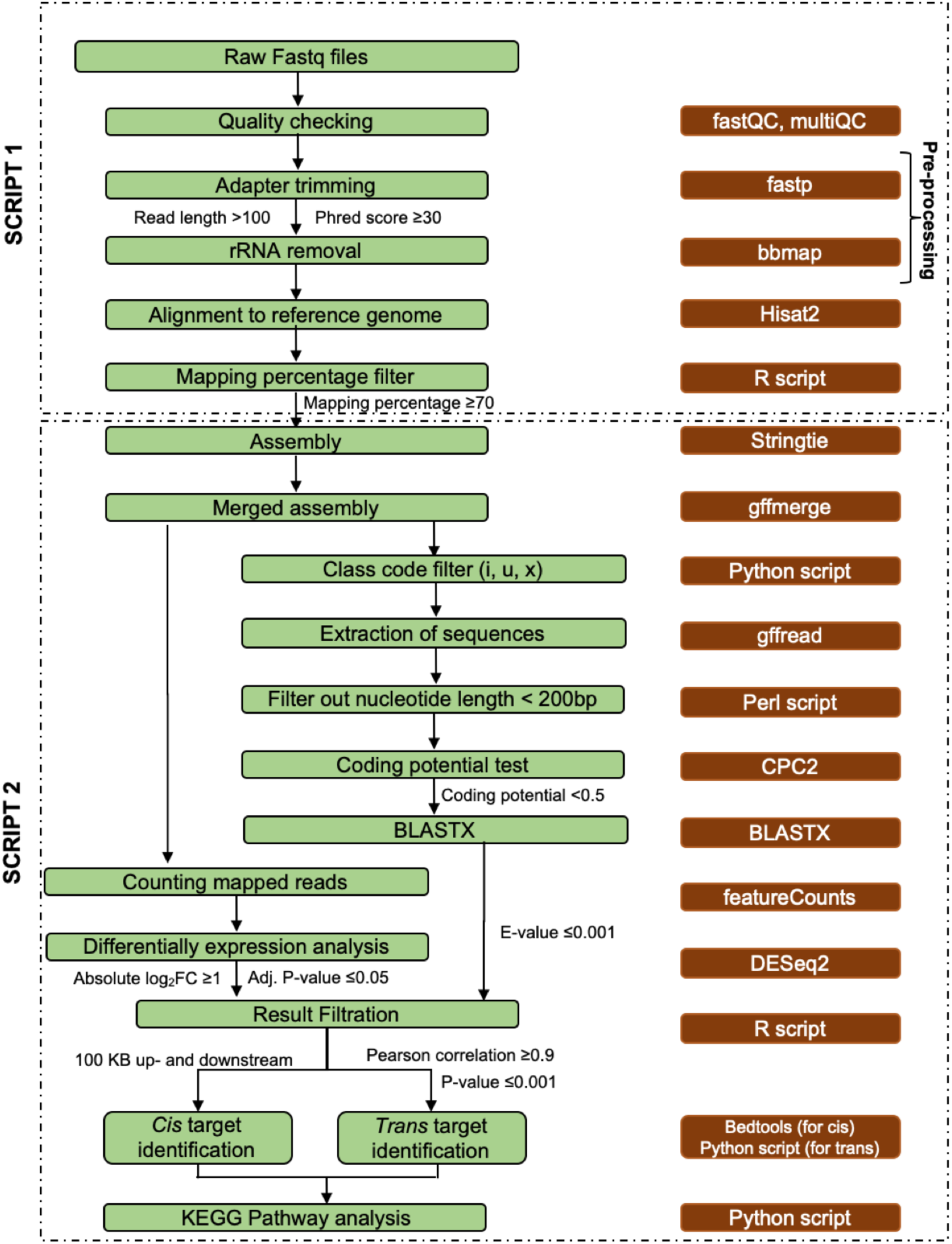
The workflow depicts the process for identification and characterization of lncRNAs. Green boxes represent the individual steps performed. Brown boxes indicate the corresponding tools used. The parameters / filters applicable are listed next to the arrows. Dashed boxes indicate the steps included in script 1 & 2. All the steps have been automated using Snakemake and can be executed with just one click.

### 2.1 Script 1- Pre-processing of the raw data and Alignment

In the initial step, all fastq files undergo quality checks using fastqc (https://www.bioinformatics.babraham.ac.uk/projects/fastqc/) and multiqc (Ewels et al. 2016). Subsequently, the fastp tool (version 0.22.0) (Chen et al. 2018) is used for quality trimming, which includes automatic detection and removal of adapters. Bases with a phred score <30 and reads shorter than 100 nucleotides are trimmed by default to enhance data quality.

Following this, the pipeline filters out any rRNA reads from the dataset. This step, achieved using bbduk (version 38.18) (https://sourceforge.net/projects/bbmap/), employs an rRNA database provided with the sortmeRNA tool (Kopylova et al. 2012). This database includes sequences from Rfam and SILVA databases. bbduk identifies and removes rRNA reads with a k-mer length of 27, retaining only those reads that do not match the rRNA database.

The preprocessed reads are then aligned to the indexed reference genome using hisat2 (version 2.2.1) (Kim et al. 2019) with the option --dta-cufflinks. This step aligns reads to the genome, generating alignment files in BAM format. Additionally, a stats file is generated that contains information about the alignment percentage. Samples with alignment percentages ≥ 70% are selected for further analysis. This ensures that only samples with high-quality alignment to the reference genome are selected for further analysis, helping to improve the accuracy and reliability of downstream analyses.

### 2.2 Script 2 - Transcriptome Assembly and lncRNA identification

The aligned BAM files, meeting the mapping cut-off criteria, are assembled using StringTie2 (version 2.2.1) (Kovaka et al. 2019). The assembly files from all samples are then merged into a single ".gtf" file using Gffutils (Pertea and Pertea 2020).

This merged file includes class code information, defining the transcript’s position in the genome. Class codes i, u, and x (i = intronic, u = intergenic, and x = antisense) are utilized to filter transcripts from the intergenic region of the genome or those not overlapping with the exonic sequence on the same strand as a pre-annotated gene. This ensures focus on the non-coding transcripts.

Using the filtered merged GTF file, a FASTA file is generated using gffread module of GFF Utilities tool (Pertea and Pertea 2020). A Perl script then filters sequences with a length ≥ 200 nucleotides, to align with the definition of lncRNA (Kapranov et al. 2007; Wang and Chekanova 2017). Each sequence then undergoes a coding potential test using CPC2 (version 1.0.1). Sequences with a coding potential ≥0.5 are eliminated as they predict for protein-coding (Kang et al. 2017). Finally, a BLASTX (Altschul et al. 1990) analysis is performed against the Swiss-Prot database (version 2.5.0) to filter out sequences similar to protein-coding genes, with the E-value parameter set to <= 0.001 by default. CPC2 calculates the coding potential based on features like ORF length, ORF integrity, isoelectric point and Fickett TESTCODE score. While BLASTX filters by cross-referencing known coding genes in the database. These double checks employed to detect for known coding genes are ideal for avoiding any false positives.

### 2.3 Script 2 - DE analysis, identification of cis and trans targets and KEGG enrichment analysis

The FeatureCounts module of the Subread tool (version 2.0.6) (Liao et al. 2014) counts the number of reads mapped to the specific co-ordinates in the chromosome and this is utilized to generate an expression count file from the BAM files. DESeq2 (version 1.40.2) (Love et al. 2014) is then employed to perform differential expression (DE) analysis, comparing the defined stages or sample types in the metadata file. The default criteria for DE analysis include a p-value cutoff of ≤ 0.05 and an absolute log2FoldChange of ≥ 1. This ensures that only genes with statistically significant and biologically meaningful changes in expression levels are considered DE, providing reliable insights into the biological processes under study.

LncRNAs regulate genes locally (*cis*) or at a distance (*trans*). To identify *cis* target genes for the differentially expressed long non-coding RNAs (DE-lncRNAs), DE genes within a 100 kb region upstream and downstream from the DE-lncRNAs are identified. This approach is based on previous studies (Tian et al. 2020; Sun et al. 2023) and is implemented using Bedtools (version 2.26.0).

For identifying *trans* targets, the DE-lncRNAs and DE genes undergo Pearson correlation analysis, with the default threshold of r2 ≥ 0.9 and a p-value ≤ 0.05. This identifies highly correlated expression patterns between DE-lncRNAs and DE-genes, helping in the identification of potential *trans* targets and offering insights into the regulatory roles of lncRNAs in gene expression.

## 3. RESULTS

### 3.1 LncRNA Identification and Classification

To demonstrate the functioning of our pipeline across multiple species we analyzed datasets from three studies associated with PRJNA657713 (Rice) (Zhang et al. 2020), GSE141035 (Sorghum) (Dhaka et al. 2020), and PRJNA1031181 (Human) (https://www.ncbi.nlm.nih.gov/geo/query/acc.cgi?acc=GSE246050). The details of the samples taken from these studies are provided in Table 1.

**Table 1.**
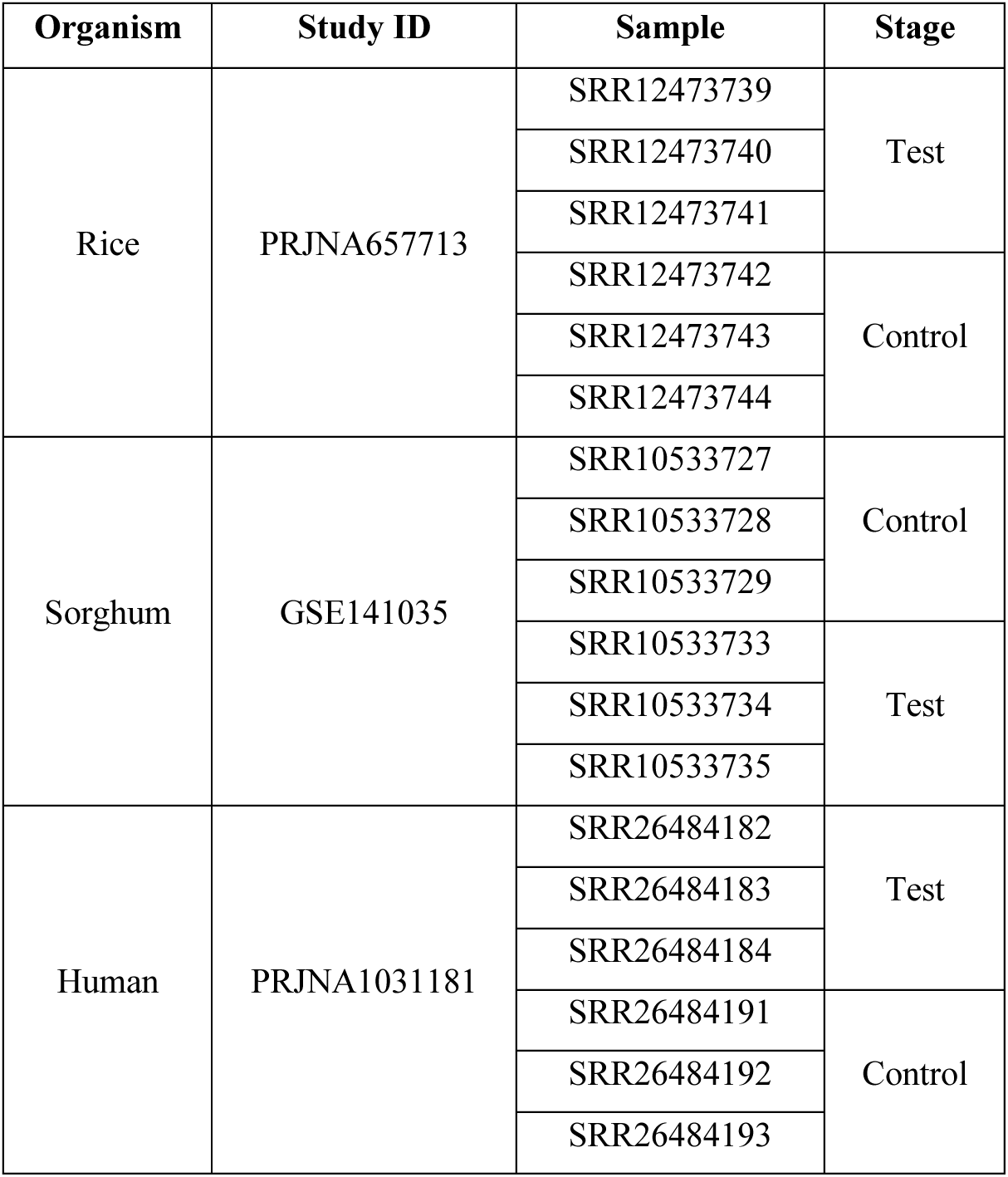
Details of different datasets used in the study.

The pipeline was run just with a single command, which parallelizes the process and optimizes it depending upon available computational resources and/or user input. Table 2, Table 3 and Table 4 summarize the overall results obtained by analyzing the three datasets. In the rice dataset (PRJNA657713), 90 DE-lncRNAs were detected. Our pipeline categorized lncRNAs into three distinct classes: intergenic, intronic, and anti-sense. Among the 90 DE- lncRNAs, 74 were intergenic, 3 were intronic, and 13 were anti-sense. Subsequently, DE-genes and DE-lncRNAs underwent analysis for cis and trans target identification. Cis targets were identified as DE-genes located within 100kb upstream and downstream regions of DE lncRNAs, leading to the identification of 53 cis targets. Pearson correlation scores were computed between the normalized read counts of DE-lncRNAs and DE-genes, resulting in the identification of 846 trans targets with a correlation coefficient (r^2) ≥ 0.9 and p-value ≤ 0.05. Out of these 53 cis and 846 trans targets, 39 were found to be both cis and trans, resulting in a total of 860 unique targets. Among these 860 targets, 110 were upregulated, and 750 were downregulated.

**Table 2.** Summary of DE-lncRNA classes identified for the analyzed datasets:

| Study ID | Organism | # of DE lncRNAs | Intergenic lncRNAs | Intronic lncRNAs | Anti-sense lncRNAs |
| --- | --- | --- | --- | --- | --- |
| PRJNA657713 | Rice | 90 | 74 | 3 | 13 |
| GSE141035 | Sorghum | 1161 | 1074 | 33 | 54 |
| PRJNA1031181 | Human | 725 | 315 | 396 | 14 |

**Table 3.** Summary of DE-lncRNAs and their associated cis and trans targets, along with enriched KEGG terms identified through the pipeline.

| Study ID | Organism | # of DE lncRNAs | # of cis targets | # of Trans Targets |
| --- | --- | --- | --- | --- |
| PRJNA657713 | Rice | 90 | 53 | 846 |
| GSE141035 | Sorghum | 1161 | 1941 | 6187 |
| PRJNA1031181 | Human | 725 | 253 | 2197 |

**Table 4.** Enriched KEGG terms identified separately for up- and down-regulated targets using the pipeline.

| Study ID | Organism | Target Upregulated | Target Downregulated | # of enriched KEGG terms |  |
| --- | --- | --- | --- | --- | --- |
|  |  |  |  | Up | Down |
| PRJNA657713 | Rice | 110 | 750 | 4 | 26 |
| GSE141035 | Sorghum | 2886 | 3317 | 18 | 16 |
| PRJNA1031181 | Human | 1163 | 1118 | 154 | 184 |

Similarly, in the Sorghum dataset (GSE141035), 1161 DE-lncRNAs were identified. Among these, 1074 were intergenic, 33 were intronic, and 54 were anti-sense. The analysis revealed 1941 cis and 6187 trans targets. Among these cis and trans targets, 1925 were identified as both cis and trans, leading to a total of 5603 unique targets. Out of these 5603 targets, 2886 were upregulated, and 3317 were downregulated.

In the human dataset (PRJNA1031181), 725 DE-lncRNAs were identified, along with 253 cis targets and 2197 trans targets. Among the DE-lncRNAs, 315 were intergenic, 396 were intronic, and 14 were anti-sense. A total of 2281 unique targets were identified, out of which 1163 were upregulated, and 1118 were downregulated.

### 3.1 Kegg Pathway Enrichment

In addition, our pipeline performs KEGG pathway enrichment analysis by applying the hypergeometric test to the up- and down-regulated target genes in each species separately. Supplementary Table 1 obtained from enrichment analysis of rice, sorghum and human lncRNA target data, displays the pathway name, gene IDs associated with each pathway, the number of genes mapped to each pathway, using a p-value cut-off of < 0.05. Using this data, the pipeline produces bubble plots in Supplementary Fig. 1 that display the statistical significance (p-value) of pathways. The plots use a colour gradient from blue to red, with blue indicating high significance and red indicating lower significance. The size of each bubble represents the number of genes associated with the pathway.

## 4. DISCUSSION

This pipeline offers the crucial benefit of processing samples simultaneously, unlike traditional methods. Our pipeline utilizes the Snakemake workflow management system to automate and run multiple tasks concurrently, streamlining the analysis workflow, reducing manual intervention, and optimizing resource utilization. It covers tasks from managing raw fastq data to conducting transcriptomics analysis, detecting and classifying lncRNAs, performing DE analysis, and identifying the *cis-* and *trans*-target genes of DE-lncRNAs. Additionally, it performs KEGG pathway enrichment analysis, providing researchers with a comprehensive understanding of lncRNA regulatory functions in cellular processes. The detailed biological application of this pipeline has been covered in one of our research article manuscript that is currently under review.

To demonstrate resource optimization and a significant reduction in processing time, we conducted a comparative analysis of manual and automated runs using Sorghum and Rice samples. Manual processing of Rice study (6 samples) required 8 hours, while Sorghum study (6 samples) took 9 hours. Utilizing our automated pipeline, the processing time decreased to 5.5 and 4.5 hours for rice and sorghum samples, respectively. This not only accelerates analysis and saves time but also enhances consistency while minimizing the risk of human error. Significantly, in our analysis, the BLASTX step stood out as the most time-consuming process, and its completion time increased in direct proportion to the sample depth.

We evaluated the pipeline’s reproducibility by conducting independent automated runs on the rice and sorghum datasets and comparing the predicted lncRNAs. We observed a high reproducibility rate of ∼ 95%, indicating that the majority of predicted lncRNAs were consistent between the runs. The remaining 5% of minor variability can be attributed to the inherent randomness in the StringTie algorithm.

During the execution of these tasks, the pipeline automatically saves the generated files and final results in the same raw data folder of the study. This approach facilitates easy data traceability, allowing users to crosscheck data from each step and aid in the identification and troubleshooting of any issues that might arise during data analysis.

Based on our literature survey we came across few tools that were designed to study lncRNA’s either in very specific contexts or to be used as analysis pipelines. We have made a list of key features needed for effective identification of lncRNA’s and other aspects highlighted earlier. The availability of these features were then compared for our pipeline and the other tools. This comparison in no way undermines the utility of those tools and their scientific rigor, as we acknowledge that some of these tools are designed for very unique scenarios and have their benefits in those scenarios. However, from the specific point of view of a comprehensive, automated, lncRNA analysis pipeline, the relevant features of these tools have been compared with our pipeline in figure 3. MechRNA (Gawronski et al. 2018) focuses solely on predicting RNA-RNA and RNA-protein interactions, requiring additional chipseq data and lacking all other features like lncRNA and its target prediction. ICAnnoLncRNA (Pronozin and Afonnikov 2023), a Snakemake pipeline, searches and classifies lncRNAs from fasta sequences specifically in plants, but does not handle raw fastq data, does not predict lncRNA targets, nor perform functional analysis. Linc2function (Ramakrishnaiah et al. 2023) offers functional annotation from lncRNA fasta sequences but does not predict lncRNAs, and it is trained on human data so it is best suited for human libraries. CALINCA (Talyan et al. 2021) specializes in identifying lncRNAs in podocyte disease, providing a podocyte-centric analysis without covering features such as target prediction and functional prediction. UClncR (Sun et al. 2017) detects lncRNAs from RNA-seq data and quantifies them but is restricted to the human genome and lacks classification, target detection, and functional prediction capabilities. lncRNADetector (Shukla et al. 2021) offers web server accessibility and user-friendliness but is specific to medicinal plant lncRNA identification without providing target and functional information. Our pipeline, focuses on lncRNA identification and analysis, operating on RNA-seq alignment files, predicting candidates, quantifying and classifying lncRNAs, predicting targets, and working across organisms. It currently cannot predict RNA-RNA or RNA-protein interactions, with efforts underway to develop this capability. Additionally, we plan to create a web-based platform for our automated pipeline for a wider reach and to enhance user-friendliness.

**Fig 3.**
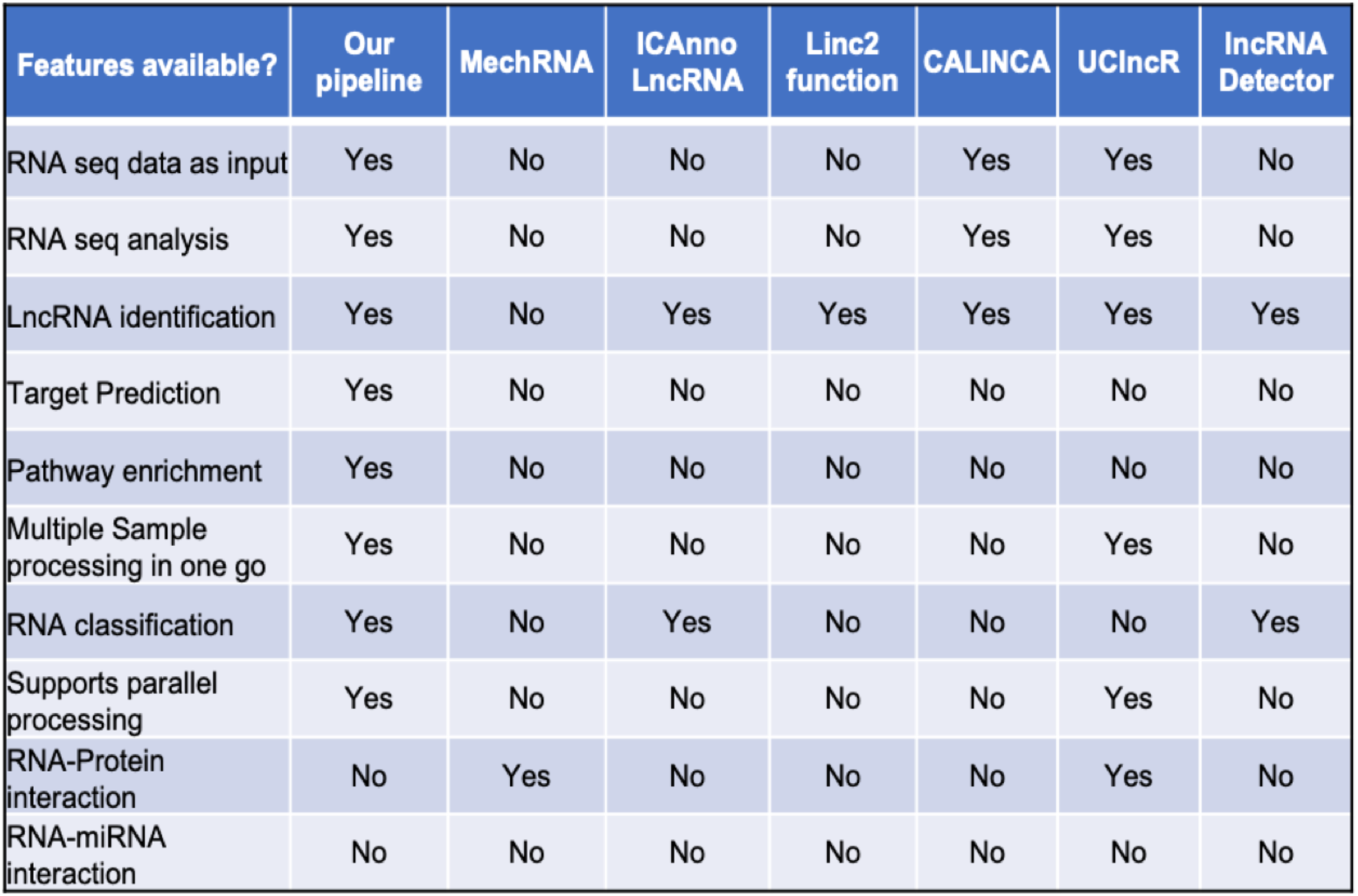
A comparative table of available tools and their features that attempt to obtain insights into lncRNA identification, classification, automation and functional aspects.

## 5. CONCLUSION

In conclusion, our SnakeMake-based pipeline provides a versatile, user-friendly, and reproducible solution for researchers seeking a comprehensive analysis of the lncRNA transcriptome. This tool will significantly contribute to advancing the field of lncRNA research by providing a standardized and efficient approach to uncovering the regulatory landscape of these intriguing molecules across different species and biological systems.

## AUTHOR CONTRIBUTIONS

MK conceived the study, CM wrote the scripts. MK and CM wrote the manuscript. BRR supervised the study, edited the manuscript for content and flow and provided critical suggestions.

## ACKNOWLEDGEMENTS

MK, CM and BRR acknowledge Jio Platforms Limited for providing the necessary facilities required and financial assistance through the salaries provided. We also acknowledge intern, Bommineni Sai, for his contributions in the script writing.

---

**Supplementary Fig. 1.**
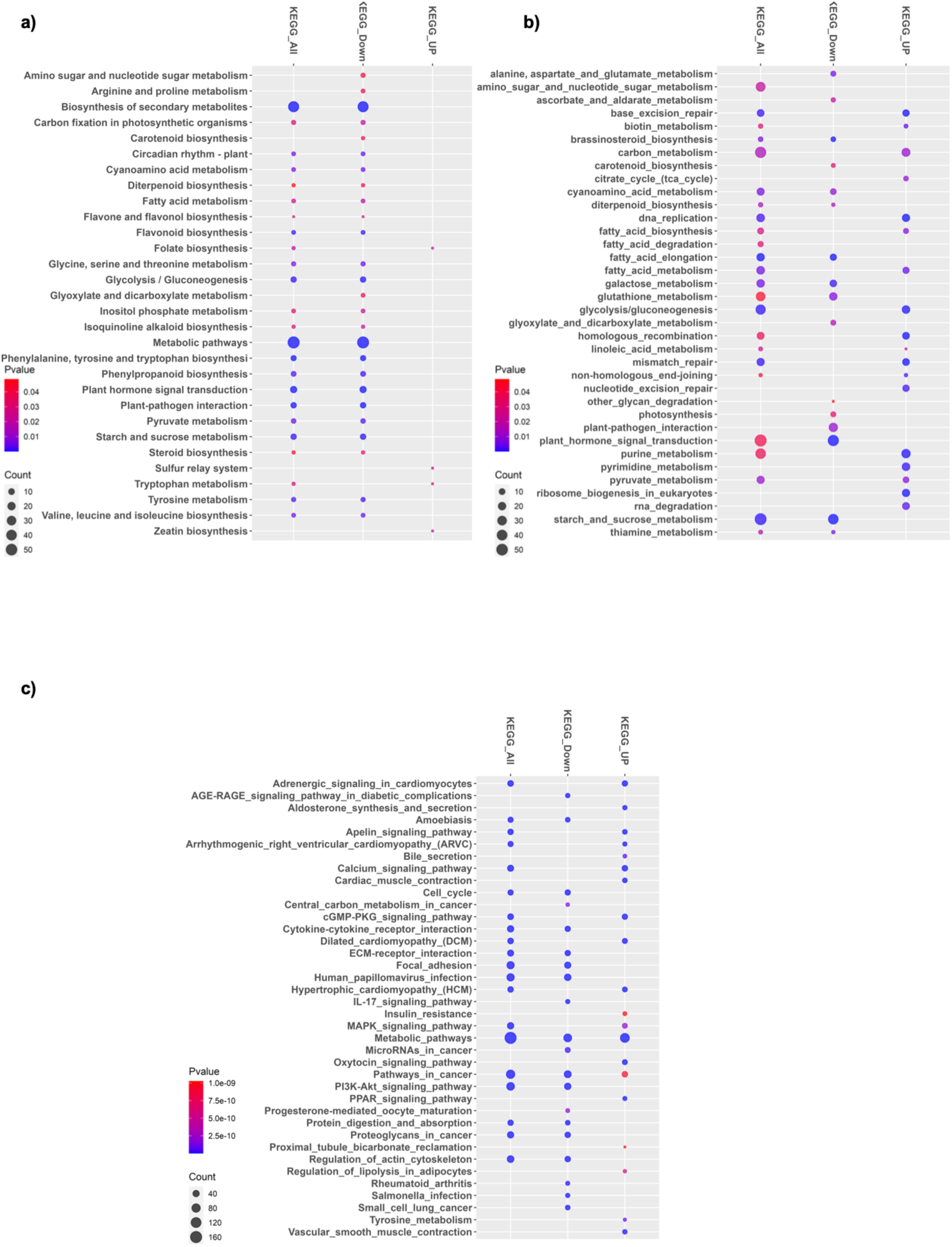
The figure displays bubble plots illustrating the enriched KEGG pathways in: a) Rice, b) Sorghum, and c) Human.

